# ORCA: Predicting replication origins in circular prokaryotic chromosomes

**DOI:** 10.1101/2024.03.28.587133

**Authors:** Zoya van Meel, Jasmijn A. Baaijens

## Abstract

The proximity of genes to the origin of replication plays a key role in replication and transcription-related processes in bacteria. Computational prediction of potential origin locations has an important role in origin discovery, critically reducing experimental costs. We present ORCA (Origin of RepliCation Assessment) as a fast and lightweight tool for the visualisation of nucleotide disparities and the prediction of the location of replication origins. ORCA uses the analysis of nucleotide disparities, *dnaA*-box regions, and target gene positions to find potential origin sites, and has a random forest classifier to predict which of these sites are likely origins. ORCA’s prediction and visualization capabilities make it a valuable *in silico* method to assist in experimental determination of replication origins. ORCA is written in Python-3.11, works on any operating system with minimal effort, and can process large databases. Full implementation details are provided in the supplementary material and the source code is freely available on GitHub: https://github.com/ZoyavanMeel/ORCA.

## 1 Background

DNA replication is an important step in the bacterial cell cycle. Most bacterial species have circular chromosomes, whose replication starts at the origin of replication (*oriC*), progresses bidirectionally, and terminates at the *terC* -region. The *oriC* -proximal distance of genes plays a significant role in the control of replication and transcription-related processes, hence determining the location of *oriC* s can further our understanding of these processes [16, 23, 25]. Moreover, *oriC* positions are also valuable in other contexts, including cross-species comparative gene location analysis. In such experiments, the *oriC* can be used as a consistent anchor point for tracking genetic shifts in lineages, the repositioning of genes, and more.

However, the *oriC* suffers from a lack of annotation, as experimental determination of the *oriC* is not straightforward. There are multiple *in vivo* and *in vitro* methods for determining the *oriC* based on protein interaction, but these are generally impractical and of low reliability. While *in vivo* methods provide native conditions for studying *oriC* -interacting proteins, these methods are restricted in their throughput and can lack quantitative assessment of the protein-*oriC* interaction. On the other hand, *in vitro* methods provide an easier way of quantifying the interaction of a protein with an *oriC*, but its results can be highly dependent on assay conditions. Because of these limitations, such methods are generally used in combination with *in silico* methods and each other [2, 12, 26].

Various tools for computationally predicting replication origins have been developed. The most widely used method for prokaryotic genomes is Ori-Finder [5, 6], an online tool which makes use of several nucleotide skew-analyses and the location of indicator genes on the chromosome. Other *oriC* -predicting tools for prokaryotes are OriLoc [7] and GenomeAtlas [11, 29]. However, OriLoc is meant mostly for visualisation and GenomeAtlas has been discontinued. Inspired by these earlier approaches, similar methods have been developed for predicting replication origins in eukaryotic genomes (e.g. Ori-Deep [22], iORI-Euk [4], OriC-ENS [1], Ori-Finder 3 [28]). However, due to the differences in replication mechanisms, such methods are unsuitable for prokaryotes [21]. Thus, to the best of our knowledge, the only currently viable option for computational prediction of *oriC* locations in prokaryotes is Ori-Finder.

Although Ori-Finder is a highly accurate tool, it is difficult to gain insights into the specific methodologies and performance because the software is not open source. Moreover, Ori-Finder is exclusively available as an online resource through a web server [5] which processes one genome at a time. Consequently, Ori-Finder is unsuitable for processing large-scale genomic datasets.

We provide ORCA (Origin of RepliCation Assessment) as an easy-to-use, open-source, and high-throughput method for predicting *oriC* s in circular bacterial chromosomes. ORCA can be fine-tuned for predictions on single organisms, but it can also be readily applied to large datasets for mass annotation. ORCA makes use of similar principles and approaches as Ori-Finder (Z-curve analysis, GC-skew, and *dnaA*(-box) locations [9, 13, 31]), with an additional model to provide confidence scores for predicted *oriC* s.

## 2 Methods & Implementation

ORCA can take as input a GenBank file (or an accession number for which it downloads the GenBank file), a genome sequence (FASTA), or a Biopython [3] SeqRecord (Figure 1A). ORCA extracts the DNA sequence and indicator gene positions for further analysis (described below), resulting in multiple candidate origins. These intermediate results can be used for plotting, or manual analysis and interpretation. We provide a Random Forest Classifier (RFC model), which takes as input the features extracted from a candidate origin and predicts a confidence score.

**Figure 1:**
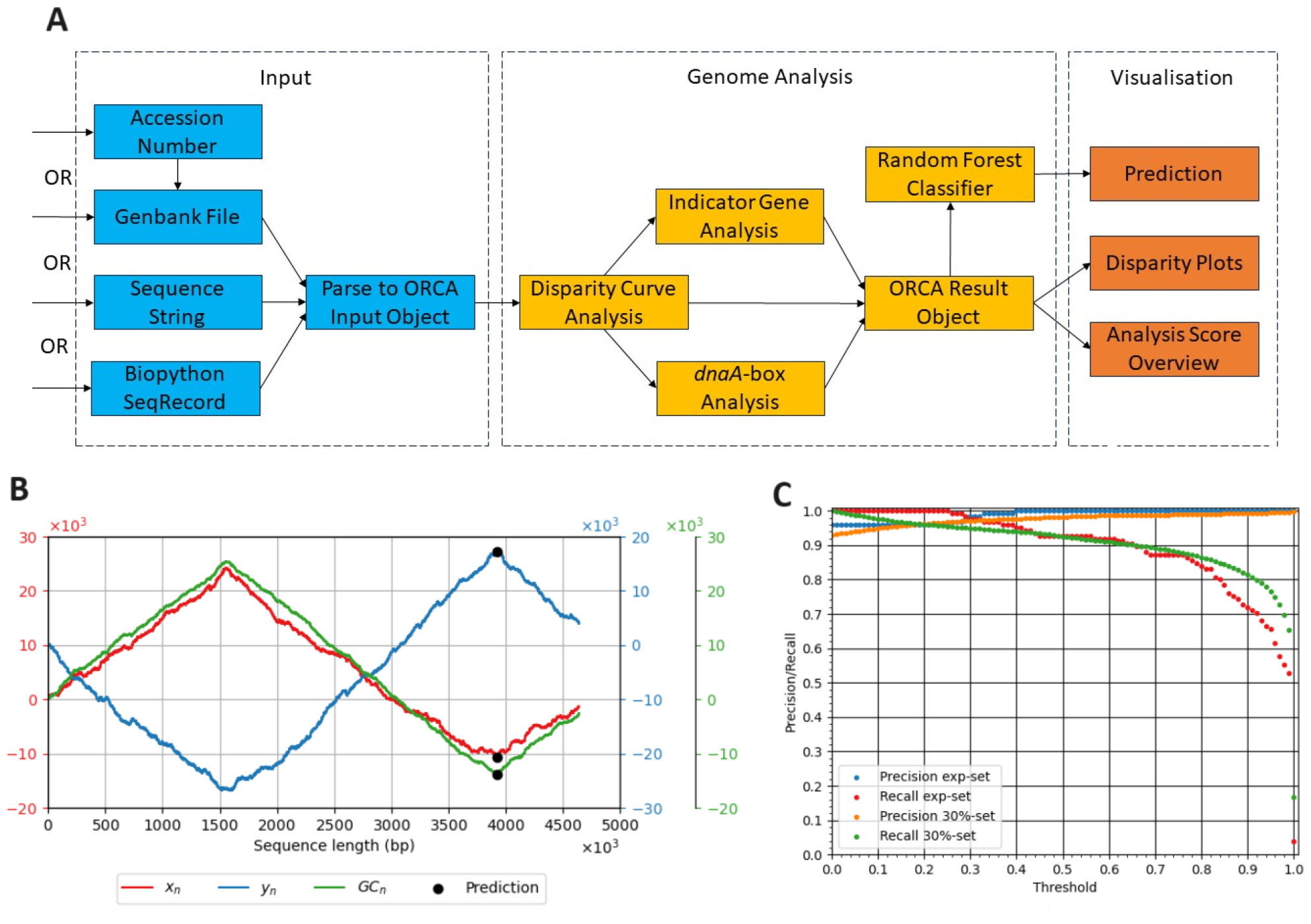
**A)** The workflow of ORCA. ORCA can handle different input types, where each is parsed accordingly to produce an ORCA input object that represents the genome sequence and indicator genes. In the middle, the genome analysis steps are shown. The Random Forest Classifier can be used for the prediction of the correctness of potential *oriC* -sites found by ORCA. **B)** An example of an ORCA plot showing the disparity curves (*x*- and *y*-components of the Z-curve, GC-skew) and the predicted origin location. This plot corresponds to *Escherichia coli* K-12 (accession: `NC_000913.3`), where the predicted origin is at 3 927 479 bp and the true origin lies at 3 925 744..3 925 975 bp [17, 19]. **C)** The precision and recall of ORCA vs. the prediction confidence threshold on both test sets. By default this threshold is set to 0.5.

### 2.1 Disparity curves analysis

The GC-disparity of a DNA sequence is defined as the distribution of guanine (G) over cytosine (C) bases. The notion of GC-disparity is generalized by the Z-Transform, an injective function that maps a DNA sequence to a 3D-curve, known as the Z-curve, by plotting three distributions of the four nucleotides against each other [30]. It was shown that minima in the *x*-component and maxima in the *y*-component of the Z-curve can be used to identify *oriC* -like regions in DNA sequences [9]. This is one of the main principles used by Ori-Finder, and we build on this for ORCA as well.

ORCA extracts positions that are characteristic to *oriC* -regions by searching for local extremes in the *x*-, *y*-, and GC-curves. It is difficult to find local extremes reliably, because there is no general size or form for the shape of the curves (see Supplementary Material). Therefore, we find extremes using SciPy [27] and keep only those which (1) represent a correct extreme (minimum or maximum) in the local neighborhood, and (2) have a matching extreme in at least one other disparity curve for the same genome. All remaining points become candidate origins and will be analyzed in later steps. We define the Z-score of a candidate origin as the number of times it was found using different neighborhood sizes. Further details on the disparity curve analysis can be found in the Supplementary Material.

### 2.2 Indicator gene analysis

For each candidate origin, we evaluate its proximity to pre-defined indicator genes, by default *dnaA* and *dnaN*. These genes are often located in close proximity to the *oriC*, since they are heavily involved in the initiation of the replication cycle [14, 18, 24]. The position of these indicator genes can help in identifying the location of an *oriC* [10]. We therefore define the G-score of a candidate origin as the average proximity to the indicator genes.

### 2.3 *dnaA*-box analysis

In parallel to the indicator gene analysis, we search for *dnaA*-boxes. These are sites where the DNA-unwinding starts during replication, with the *dnaA*-protein binding to *dnaA*-boxes [8, 15]. Hence, the more *dnaA*-boxes are close to a candidate origin, the more likely this site is to be a true origin. The *dnaA*-boxes are characterized by sequences of nine bases, for which ORCA uses the base sequence TTATNCACA as default [20]. We assign to each candidate *oriC* a D-score, which counts how many *dnaA*-boxes are found in its local genomic neighbourhood. By default this neighborhood consists of a window whose length equals 5 % of the total chromosome size, centered on the candidate site.

### 2.4 Final prediction using a Random Forest Classifier

ORCA computes Z-, G-, and D-scores for all candidate origins and outputs an object that can be used with the provided plotters to visualise the Z-curve, GC-skew, and the predicted origin(s). For the final prediction, ORCA provides an RFC model that was trained to predict a confidence score for each candidate origin, using the Z-, G-, and D-scores as features. For each chromosome the candidate origin with the highest confidence scores is selected to be the final prediction.

## 3 Results & Discussion

For training and testing, we used a dataset consisting of all accessions present in the complete and chromosome-level assemblies for circular genomes in the DoriC 12.0 database [6], downloaded on May 16th, 2023. The dataset was split into one training set and two testing sets. The first testing set (“exp-set”) consisted of 32 organisms, comprising 37 origins in total, which were experimentally verified in previous studies (see Supplementary Material). The remaining data was divided in a 70:30 stratified train:test split. This test set is referred to as the 30%-set. Since the features are calculated independent of the homology of the organism, splitting was performed randomly. Further details on the model training and validation can be found in the Supplementary Material.

We evaluate ORCA in terms of precision and recall. The RFC model provides confidence scores for its predictions, which can be used as a confidence threshold. We ran ORCA for each sample in the dataset, and considered the candidate *oriC* with the highest confidence score as the final prediction. A prediction is considered a true positive when it is located at most 2 % of the chromosome length away from the midpoint of the true origin. Note that the DoriC database was built using Ori-Finder [5], so by assessing performance on DoriC we are comparing ORCA to Ori-Finder.

Figure 1C shows precision and recall for confidence thresholds ranging from 0.0 to 1.0. We observe a natural precision-recall trade-off: with increasing confidence thresholds, precision increases while recall decreases. At a threshold of 0.50, ORCA achieves a precision/recall of 0.98/0.93 on the 30%-set and 1.00/0.93 on the exp-set.

With ORCA we provide an easy-to-use tool for the prediction and analysis of the location of origins of replication in circular prokaryotic DNA. ORCA is highly adaptable and can be tuned towards specific use-cases. For example, the target genes used in the G-score can be made specific to the species of interest. The implementation is lightweight, making it ideal for high-throughput research, and can easily be integrated into genome analysis pipelines.

## Data availability

The data underlying this article are available on Zenodo (https://dx.doi.org/10.5281/zenodo.10888580) as well as an archived version of the code (`10.5281/zenodo.10888686`) to fully reproduce the results.

## Author contributions

Z.v.M. conceived and conducted the experiments. All authors analysed the results, wrote and reviewed the manuscript.

## Notes

https://github.com/ZoyavanMeel/ORCA

https://zenodo.org/records/10888687

https://zenodo.org/records/10888581

### Competing Interest Statement

The authors have declared no competing interest.

