## Supplementary Material for "ORCA: Predicting replication origins in circular prokaryotic chromosomes"

### ORCA: Predicting replication origins in circular prokaryotic chromosomes
### Supplementary Material

Zoya van Meel[1] Jasmijn A. Baaijens[1,2]

[1]Delft University of Technology, Delft, Netherlands
[2]Harvard Medical School, Boston, USA

ORCA's workflow happens in several steps, outlined in Figure 1 of the main paper. This supplement explains the algorithmic details of the genome analysis.

#### 1 Disparity curve analysis

The first step is the disparity curve analysis. ORCA first maps the full Z-curve and GC-skew of the given sequence and makes a *dnaA*-box dictionary for the positions of each of the provided *dnaA*-boxes. All further analyses are done on these curves and dictionary. The sequence is still returned in the ORCA result object, but is no longer used in the computation. The disparity curve analysis is done on the $x$-, and $y$-components of the Z-curve as well as the GC-skew curve. These three curves are analysed for candidate origins and these candidates are used in the indicator gene and *dnaA*-box analysis later in the workflow.

First, SciPy [1] is used to detect local extremes in each of the three aforementioned curves. In this step, the curves are treated as random signals. From visual inspection of plotted disparity curves, no generalised form of the Z-curve could be derived. It could be possible to optimize this peak detection problem for specific species, since closely related species will likely produce similar shaped disparity curves, but not across all prokaryotes with circular DNA. There is not only the issue of DNA recombination across species and organisms, but also the issue of linearisation. Because the DNA is circular, there is no definite place to start the linearised sequence. Starting the linearisation of a circular chromosome at the origin of replication could ensure some consistency in future analyses, but that is not something this project could benefit from. There is an informal convention to publish the linearised sequences of circular chromosomes starting at the *oriC*, but it is never noted whether this is done or not.

These factors makes the detection of local extremes very difficult. It is possible that there are large clusters of sequences that produce similar shaped disparity curves that only look dissimilar because of linearisation. However, without manually checking each of them, it is not possible to know which those are. Figure 1 shows examples of the disparity curves of various chromosomes in the dataset. It can be seen that generalising these curves is not easily possible.

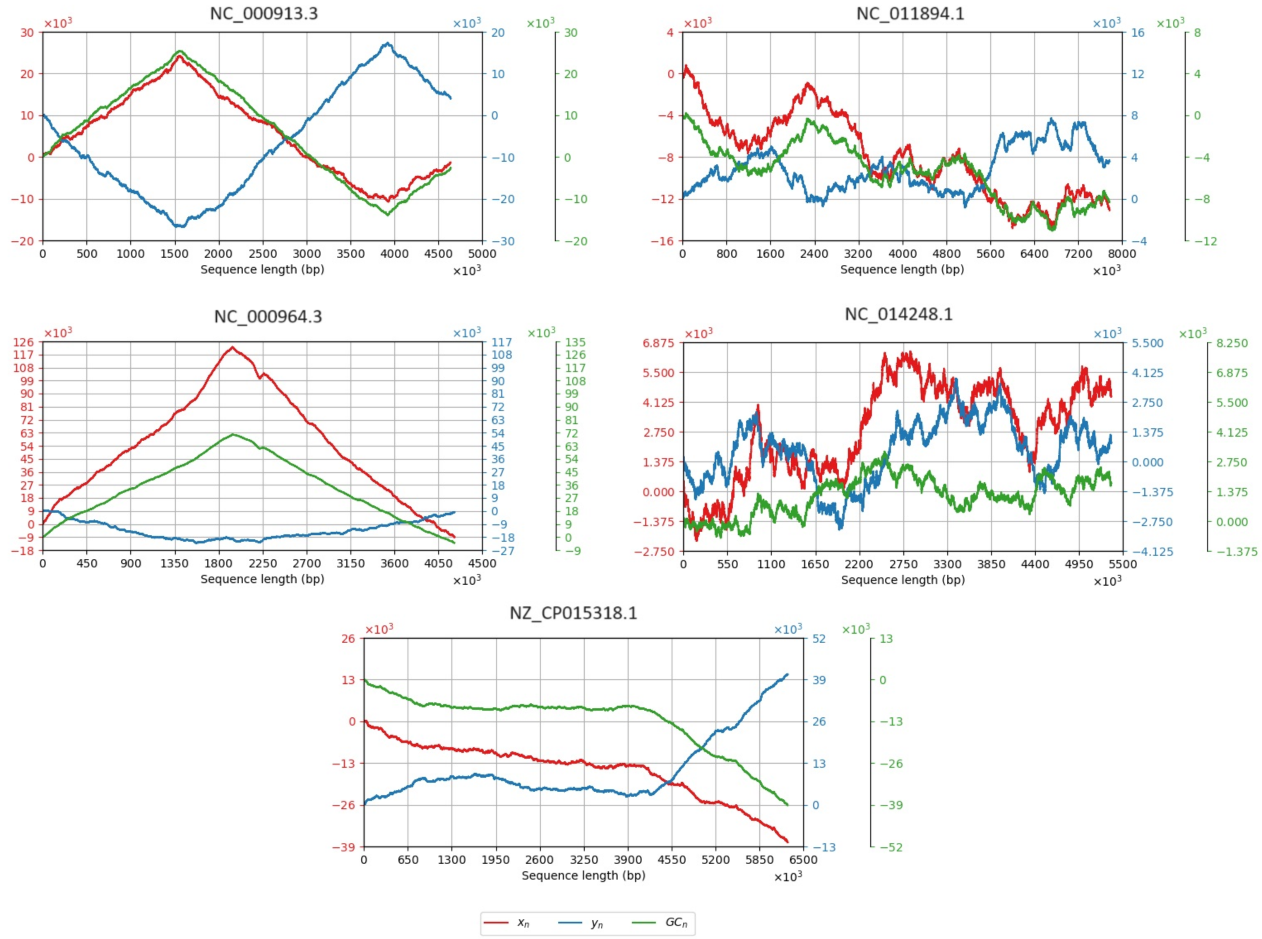


**Figure 1:** An overview the disparity curves of different organisms. From upper left to lower right: `NC_000913.3` = *Escherichia coli* K-12; `NC_011894.1` = *Methylobacterium nodulans* ORS 2060; `NC_000964.3` = *Bacillus subtilis* 168; `NC_014248.1` = *Nostoc azollae* 0708; `NZ_CP015318.1` = *Mesorhizobium amorphae* CCNWGS0123.

Figure 1 shows a few different examples of the disparity curves of various chromosomes in the dataset. It can be seen that generalising these curves is not easily possible. The sequence of `NC_000964.3` has been linearised on the *oriC*. Here it is clear that the *oriC* is found in the minima of the $x$-component and GC-skew and the maximum of the $y$-component. The same is true for `NC_000913.3`, but this sequence is not linearised on the *oriC*. If this sequence had been linearised on the *oriC* as well, we would have curves with a similar shape to `NC_000964.3`. However, without prior inspection, it is not possible to know that these curves are similar. One way to reduce the effects of linearisation could be to run the analysis with varying shifts in the sequence. This was an approach used by Oriloc for visualisation [2].

Figure 1 also shows the disparity curves of three other bacteria. It can be seen that these differ much more from both the previous two sequences and each other. This shows that linearisation is not the only issue in peak detection. Therefore, our approach to peak selection involved an initial selection of an large pool of candidates followed by a filtering process. Using SciPy, we look for both maxima and minima in each curve that are distanced around 1/12th of the length of the sequence apart. We also add the global extremes. This gives us at most 26 unique candidate origins to be filtered and analysed.

#### 1.1 Candidate Filtering

Filtering the candidate pool is a two-step process. Both filters are run on all extremes per curve. An extreme is rejected for further examination if it gets rejected by either filter. First, a filter checks if the point is truly an extreme in its own window. The type of extreme is specified for each curve. Since both maxima and minima are considered at the start, this filter will eliminate all minima when looking for maxima, and vice versa. This filter represents a check if any given point is the wrong type of extreme or if the point is on a slope that was not caught by SciPy. For example, the $x$-component of `NC_000964.3` in Figure 1 shows a local minimum just under 2.25 Mbp. Depending on the window size, this extreme will pass the filter, being considered a true local minimum, or it will be rejected for being on a slope.

The second filter checks the extremes against each other. All combinations of point pairs are compared against each other. If the pair does not have an intersecting window, they are ignored. If their windows do intersect, then the points are checked for which is a more desirable extreme out of the two. For example, if one is looking for maxima, then the smaller one of the two is rejected.

#### 1.2 Curve Combinations

This step aims to line up the accepted extreme points from the previous step to each other. This step happens per specified window size. Since there are three curves that are being analysed, three groups of accepted extremes have to be lined up to each other. Finding an extreme in one curve does not mean much if there is no corresponding point around the same position on another curve. In this step we check all pairs of curves (($x$, $y$), ($x$, GC), ($y$, GC)) and see if the candidates found from each curve have a counterpart in another curve. For this, we check if any points from one curve intersect their window with any points from another curve. All candidates that found a match, get turned into a candidate at the centre of both points.

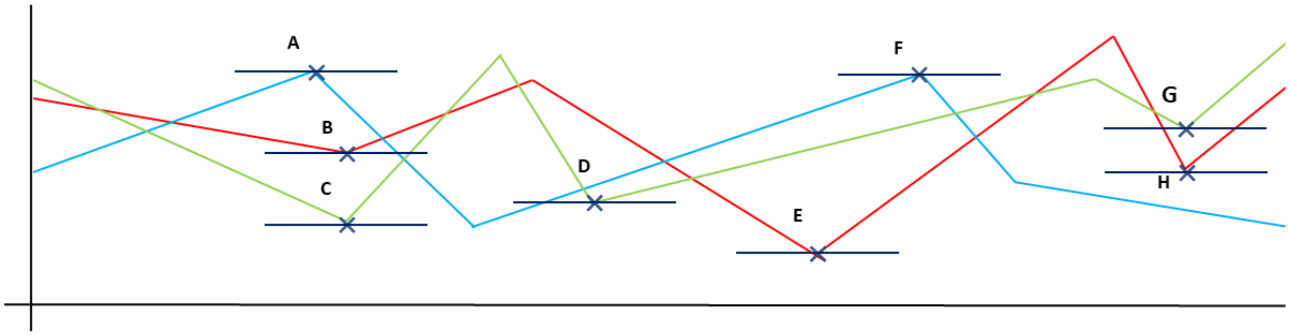


**Figure 2:** An example of how ORCA would match the given extremes shown in the plot. This disparity plot is for illustrative purposes only and was not generated from any genome. The green line represents the GC-skew and the red and blue lines represent the $x$-, and $y$-components of the Z-curve, respectively. The labelled crosses represent local extremes found on each curve. The horizontal lines trough them are their respective windows.

An example of this process is shown in Figure 2. This example disparity plot shows how different groups of local extremes could be merged into candidate origins. The first group to examine include the points A, B, and C. These extremes do not line up perfectly, but, depending on the window size, these points will still be considered as representing the same significant change in the chemical properties of the sequence. A, B, and C will be paired at each iteration.

Next, we examine point D. This point does not intersect its window with any other point on any of the other curves. Since this extreme is not represented on any other curve, we do not consider it for any further analysis.

Points E and F may seem far apart, but their windows still intersect and will therefore be considered as representing a distinct change in the sequence. This change is only found in two of the three curves, but will still be considered for further analysis. The same is true for points G and H.

Now we have a group of merged candidate origins that appear in at least two out of the three curves. These candidates will be considered as being present on a sufficient amount of curves. We now wish to merge any clusters of candidates. For example, imagine there are three candidates, all sufficiently close to each other that they were matched. This means there is one replacement candidate for each match:

- Point P from A and B.
- Point Q from A and C.
- Point R from B and C.
- Point S from E and F.
- Point T from G and H.

These five points would continue to the next step in the process. Points P, Q, and R would all be representing a match between the original points A, B, and C. The reason we do not match A, B, and C immediately is that doing the matching in pairs gives more leeway to the matches that can be made. It also makes it so that extremes found in similar locations across all disparity curves are scored higher on average in the Z-score calculation. A group of three original extremes will still be represented by three matches after this step, but a match between two extremes that is only present in two curves, will be represented by only one new point.

##### 1.3 Connected Components

We also ran the previous two steps for three different window sizes (1, 3, and 5 % of the chromosome length as default). This leaves us with a lot of points around roughly the same positions. Take the the previous points P, Q, and R as an example. These points have to be clustered together. There is also the possibility that similar points as these were found when using a different window size. We cluster these candidates by representing the clustering as a graph problem. We try to find connected components in an undirected graph. Figure 3 shows a simplified example of this step. Since we have found and matched all points that appear in the same positions across all curves, we can imagine them in the same one dimensional space using the indices of the points on the sequence. The threshold for any components being connected is set by the `max_point_spread` parameter and is set to 5 % of the chromosome length as default.

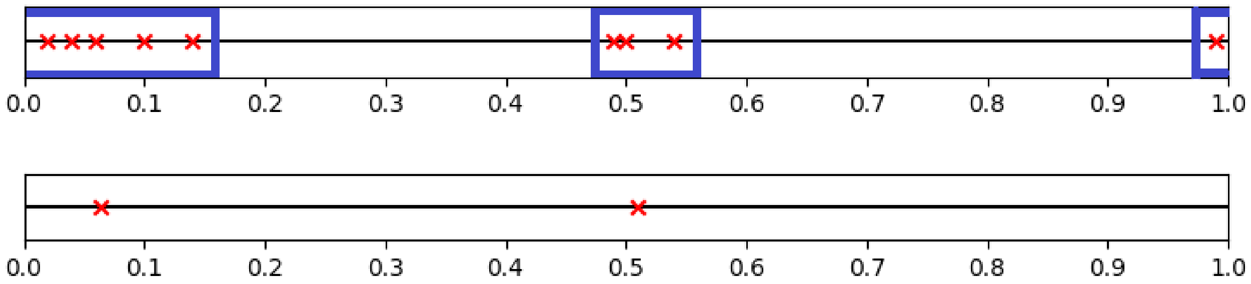


**Figure 3:** Example illustration of the intended effect of merging connected components. The top picture shows two clusters of points in a 1D space representing a simplification of a disparity curve. The boxes surrounding the points represent the groups that will be merged into one point. The bottom picture shows the result of the merger.

It is technically possible to chain all points together, provided each point falls within threshold distance of another point already in the connected component. This is why we set the maximum within-distance of each connected component to 1.5 times the threshold. The steps below attempt to outline the self-regulation of this algorithm:

1. Find connected components making use of the `max_point_spread` as threshold $t$.
2. We keep track of which threshold $t_u$ was used when connecting each group.
3. Check the within-distance $d_w$ of each group. If $d_w > 1.5 \cdot t$: flag the group for reconnection.
4. If the group is flagged, we rerun the connected components algorithm on only the points in this group. This time we use threshold $t_u \leftarrow 0.75 \cdot t_u$.
5. We continue to run a flagged group with smaller thresholds until the within-distance is accepted or $t_u < 0.25 \cdot t$. If $t_u$ is this small, we simply accept the group even if the within-distance is too large.

##### 1.4 Z-score

Before we merge the connected components as illustrated in Figure 3, we use the number of elements in the groups in calculating the Z-score. For that we use the following formula:

$$\mathrm{Z}_i = \frac{\text{length of group}_i}{\text{total number of candidates}}$$

for $i = 0, 1, 2, ..., n-1$ with $n$ as the number of groups found by the connected components algorithm. We calculate the centre of each group. Those points are the candidate origin points that are used in the score calculations. These candidates are presented to the user. Performing all previous steps leave 93.3 % of all tested samples with five or fewer candidate origins and 99.9 % of samples with ten or fewer candidates for scoring.

#### 2 Indicator gene analysis

Compared to the disparity curve analysis, the indicator gene analysis is very simple. We continue with the candidates computed in the previous step. We calculate a distance matrix of each candidate against each indicator gene (*dnaA* and *dnaN* as default). We scale this matrix between 0 and 1 inclusive. Then we take the average distance of each candidate to the indicator genes and label this the G-score. This can be formalised as follows:

$$\mathrm{G}_i = \frac{\sum_j^k d_{i,j}}{k}$$

where $d_{i,j}$ is the scaled distance of candidate $i$ to gene $j$ and $k$ is the number of indicator genes. If no indicator genes were found or specified, the G-score of each candidate is equal to zero.

### 3 *dnaA*-box analysis

The *dnaA*-box analysis is also performed per candidate. We make a positional dictionary of each allowed *dnaA*-box. Allowed *dnaA*-boxes are the set of all *dnaA*-boxes provided (TTATNCACA as default [3]) and all boxes with the allowed number of mismatches (0 as default). All boxes are weighted equally. *dnaA*-boxes that have no mismatches with the provided boxes are not favoured over those that do have mismatches. The D-score can be formalised as follows:

$$\mathrm{D}_i = \frac{\text{number of } dnaA\text{-boxes contained by } i}{\text{total number of } dnaA\text{-boxes present in all candidates}}$$

where candidate $i$ contains *dnaA*-box $j$ if the distance $d_{i,j}$ is smaller than half the size of the largest provided window size (5 % of the chromosome length as default). If no candidates contained *dnaA*-boxes, then the D-score for each is zero.

### 4 Random Forest Classifier

Each candidate has a Z-, G-, and D-score. These scores can be manually analysed by the user along with visual inspection of the relevant disparity curves or they can be used as features for the provided the Random Forest Classifier (RFC). Since all scores have already been scaled between 0 and 1 inclusive, no further preprocessing of the features is necessary. The provided RFC has been trained on all 24 thousand entries in DoriC 12.0 using ORCA's default parameters. If there is any deviation from the standard parameters, the performance of the RFC can no longer be guaranteed and retraining is recommended.

The model was tuned and validated using Scikit-learn's `GridSearchCV` class [4]. This class performs a gridsearch over the provided hyperparameters and selects on the parameters that give the best performance. Best performance is defined as the highest average score over the individual cross-validation sets. The scores that were used were precision and recall. The cross-validation was 5-fold and stratified, implemented using Scikit-learn.

Note that 13.0 % of the chromosomes in the DoriC dataset (13.5 % of the exp-set) have a bipartite origin, meaning that two locations within close proximity have been annotated as origins of replication. Current putative bipartite origins are separated by at most 150 kilobases, and the potential reasons for the existence of bipartite origins are numerous and speculative [5]. Therefore, in case of a bipartite origin annotation, we consider the predicted origin a true positive if it lies at most 2 % of the genome size away from at least one of these origins.

### 5 Experimentally verified *oriC*s

**Table 1:** Overview of the experimentally verified *oriC* positions. The results from each of the papers were cross-referenced with the corresponding available sequences to pin down the exact starting and ending bases for each origin.

| Accession number | Starting base | Ending base | Source |
|---|---:|---:|---|
| NC_000913.3 | 3925744 | 3925975 | Oka, Sugimoto, Takanami, *et al.* [6] |
| NC_002947.4 | 8947 | 9541 | Yee and Smith [7] |
| NC_000964.3 | 4214414 | 409 | Moriya, Tsujikawa, Hassan, *et al.* [8] |
| NC_000964.3 | 1748 | 1938 | Moriya, Tsujikawa, Hassan, *et al.* [8] |
| NC_007633.1 | 1009729 | 1010023 | Fujita, Yoshikawa, and Ogasawara [9] |
| NZ_CP050522.1 | 4269844 | 4270780 | Calcutt and Schmidt [10] |
| NC_002971.4 | 1835664 | 1836065 | Suhan, Chen, Thompson, *et al.* [11] |
| NC_000962.3 | 1525 | 2051 | Salazar, Fsihi, Rossi, *et al.* [12] |
| NZ_CP054795.1 | 5902438 | 5903030 | Salazar, Fsihi, Rossi, *et al.* [12] |
| NZ_CP029543.1 | 1567 | 2080 | Salazar, Fsihi, Rossi, *et al.* [12] |
| NC_006461.1 | 1849496 | 54 | Schaper, Nardmann, Lüder, *et al.* [13] |
| NC_000117.1 | 719988 | 720258 | Stephens, Kalman, Lammel, *et al.* [14] |
| NZ_CP011330.1 | 1141221 | 1141374 | Zawilak, Cebrat, Mackiewicz, *et al.* [15] |
| NC_002696.2 | 4016703 | 234 | Marczynski and Shapiro [16] |
| NC_020528.1 | 73 | 477 | Sibley, MacLellan, and Finan [17] |
| NC_003047.1 | 73 | 477 | Sibley, MacLellan, and Finan [17] |
| NC_011916.1 | 4042684 | 201 | Taylor, Ouimet, Wargachuk, *et al.* [18] |
| NZ_CP009467.1 | 209217 | 209684 | Zyskind, Cleary, Brusilow, *et al.* [19] |
| NC_003197.2 | 4083801 | 4084177 | Zyskind and Smith [20] |
| NZ_CP041925.1 | 5147608 | 5147989 | Cleary, Smith, Harding, *et al.* [21] |
| NC_016845.1 | 1 | 381 | Cleary, Smith, Harding, *et al.* [21] |
| NZ_CP051652.1 | 4840535 | 4840923 | Takeda, Harding, Smith, *et al.* [22] |
| NC_005363.1 | 1417 | 1648 | Makowski, Donczew, Weigel, *et al.* [23] |
| NC_019567.1 | 1417 | 1648 | Makowski, Donczew, Weigel, *et al.* [23] |
| NC_018939.1 | 1607419 | 1607646 | Donczew, Weigel, Lurz, *et al.* [24] |
| NC_018939.1 | 1609021 | 1609173 | Donczew, Weigel, Lurz, *et al.* [24] |
| NC_009850.1 | 2341023 | 2341251 | Jaworski, Donczew, Mielke, *et al.* [25] |
| NC_009850.1 | 1318 | 1474 | Jaworski, Donczew, Mielke, *et al.* [25] |
| NC_007575.1 | 2201475 | 14 | Jaworski, Donczew, Mielke, *et al.* [25] |
| NC_007575.1 | 1323 | 1490 | Jaworski, Donczew, Mielke, *et al.* [25] |
| NC_005090.1 | 2110228 | 2110355 | Jaworski, Donczew, Mielke, *et al.* [25] |
| NC_005090.1 | 1315 | 1473 | Jaworski, Donczew, Mielke, *et al.* [25] |
| NC_010546.1 | 1886587 | 1887114 | Huang, Song, Yang, *et al.* [26] |
| NC_003869.1 | 2689326 | 365 | Pei, Liu, Li, *et al.* [27] |
| NC_003272.1 | 2404398 | 2404811 | Zhou, Chen, Wang, *et al.* [28] |
| NC_007604.1 | 2695871 | 109 | Watanabe, Ohbayashi, Shiwa, *et al.* [29] |
| NC_008255.1 | 4052571 | 4053394 | Xu, Ji, Chen, *et al.* [30] |
